# Macrophages migrate persistently and directionally upon entering 2D confinement in the presence of extracellular matrix

**DOI:** 10.1101/2025.05.15.654321

**Authors:** Matthew W. Stinson, Summer G. Paulson, Ethan M. Carlile, Jeremy D. Rotty

## Abstract

Cells sense and respond to their environment in a myriad of ways. In many instances they must integrate simultaneous cues ranging from the physical properties and composition of the extracellular matrix to guidance cues that stimulate chemotaxis or haptotaxis. How cells make sense of multiple simultaneous cues is an ongoing physiologically relevant question. The present study seeks to contribute to the understanding of multi-cue sensing by understanding how the transition to a confined setting with or without an added haptotactic gradient alters macrophage migration. We found that the transition to confinement is itself a directional cue capable of driving persistent migration hours after macrophages enter the confined environment. Next, we found that a haptotactic fibronectin gradient made cells even more directionally persistent under confinement. Finally, Arp2/3 complex deletion rendered macrophages unresponsive to the haptotactic gradient, but they retained directionally persistent migration due to their transition to confinement. These findings may be particularly relevant for cells that move from an adherent 2D environment into a confining 3D environment, like leukocytes and circulating tumor cells that extravasate into peripheral tissue.

**Summary Statement:** Macrophages migrate persistently after they transition from 2D adhesion to an adhesive confined environment. Migration is enhanced further if cells sense a fibronectin gradient. This may have relevance to immune and cancer cell behavior.

## ITRODUCTION

Integration of simultaneous cues in the microenvironment is a fundamental problem that cells need to solve in many *in vivo* contexts. Cells respond to soluble or substrate-bound cues, changes in substrate stiffness or topology, and electric fields (SenGupta et al., 2021). Other powerful cellular guidance cues include hydraulic pressure (Lennon-Dumenil and Moreau, 2021; Moreau et al., 2019), extracellular viscosity (Bera et al., 2022), and following the path of least resistance through a 3D environment (Renkawitz et al., 2019), among many others. While much has been learned about each of these guidance systems, the *in vivo* environment is complex and understanding how these diverse factors interact with one another requires more investigation.

Many cells sense and respond to 3D confining environments, and there are increasingly sophisticated strategies in place to tune the characteristics of these assays *in vitro* (Kameritsch and Renkawitz, 2020). Confinement devices, microchannels, pillar forests, under-agarose assays, and many others have been used in recent years to interrogate cellular decision making, often in the context of chemotactic migration (Kameritsch and Renkawitz, 2020). However, other elements of the complex microenvironment have been less widely interrogated. One such factor is how the transition to a confined setting may inherently influence cellular responses like motility. Leukocytes and cancer cells broadly share the ability to extravasate from a 2D environment in the vasculature into confining 3D tissue environments (Mondadori et al., 2020; Strilic and Offermanns, 2017). It remains unclear whether this transition itself can be sufficient to drive further directional penetration into peripheral tissue.

In addition, the contribution of extracellular matrix sensing (ECM) to cellular responses is less well-defined than chemotaxis, especially under confinement. Haptotaxis refers to a cell’s ability to sense and migrate toward an increasing concentration of surface-bound directional cues, like the glycoproteins that compose the ECM (Carter, 1965). Neurons (Varadarajan et al., 2017), dendritic cells (Weber et al., 2013), macrophages (Rotty et al., 2017), platelets (Nicolai et al., 2020), fibroblasts (King et al., 2016), and non-adherent T-cells (Luo et al., 2020) all haptosense. Adhesion (Liu et al., 2015) and haptotactic sensing (Weber et al., 2013) influence migration in confined settings, and haptotaxis has been linked to cancer (Oudin et al., 2016), fibrosis (Wen et al., 2015), wound healing (Sawicka et al., 2015), and inflammation (Nicolai et al., 2020). Integrins activate and cluster at higher substrate concentrations, leading to enhanced recruitment of adhesive effectors, such as talin and vinculin, and initiation of migratory signaling (e.g. Src and FAK) at the leading edge (King et al., 2016; SenGupta et al., 2021; Wu et al., 2012). The adhesive and signaling cues converge on the branched actin-nucleating Arp2/3 complex, which contains seven subunits: the actin related proteins Arp2 and Arp3, and Arpc1-5 (Rotty et al., 2013). Branched actin polymerization drives leading edge lamellipodial protrusion in haptotactic contexts, as demonstrated by work conducted in fibroblasts, macrophages, and platelets (Nicolai et al., 2020; Rotty et al., 2017; Wu et al., 2012). On the other hand, integrin-based ECM adhesion seems to be less important for interstitial directed migration in some *in vivo* contexts (Lammermann et al., 2008). Therefore, it remains unclear how and in which contexts ECM cues collaborate with confinement to direct cellular behavior.

We used the version of the classic under-agarose assay (Nelson et al., 1975) described by Heit and Kubes (Heit and Kubes, 2003) to demonstrate that macrophages move radially away from their starting location in a directionally persistent fashion after entering confinement. Fibronectin gradients enhanced macrophage directional migration after entering confinement even further than confinement alone. Arp2/3 disruption removed the influence of the fibronectin gradient, and lowered macrophage directional persistence compared to WT counterparts that still haptotax. Arp2/3 KO macrophages respond to confinement with an overall positive FMI and high persistence, indicating that the Arp2/3 complex is not required for confinement-induced radial migration. The transition to confinement may be a directional cue for myeloid cells that is reinforced by the local ECM upon entering a target tissue. These findings may be broadly relevant for leukocytes and circulating tumor cells that move into confined 3D environments from the peripheral circulation.

## RESULTS

### Macrophages move directionally upon entering confinement

Our under-agarose confinement method uses wells punched in a 1% agarose gel polymerized in a glass-bottom dish (**Fig. 1A**). The middle well here is referred to as the ‘source’ where media or a directional cue (e.g. fibronectin) is applied to the system (see **Fig. 1A**). The ‘far’ distance of mm is based on Heit and Kubes (Heit and Kubes, 2003). We also chose to make a ‘close’ well at 1.25 mm in case different directional cues require distinct gradient conditions. Two zones were established as cells moved under the agarose over 16 hours. As depicted, Zone 1 encompasses the edge of the cell well and the area directly outside of it. Zone 2 begins at the edge of Zone 1. We focused our quantitative analyses on cells in Zone 2. Cells in this zone move independently and do not often physically interact with cells around them, which is a complicating factor that can artificially alter migratory characteristics. Red gradient symbols in this schematic represent the relative strength and orientation of the fibronectin gradient (when present). Finally, (+) and (-) designations correspond to cells entering agarose in the direction toward the center well (+) or away from it (-). Fluorescent dextran-infused agarose gels clearly demonstrate that cells were encapsulated under agarose using this system. Dextran signal was detected surrounding cells and within them, indicating that macrophages directly interacted with and internalized the dextran in the gel (**Fig. S1A**). Additional evidence of confinement comes from the noticeable filamentous actin (F-actin) puncta generated by macrophages under agarose (**Fig. S1B**), similar to structures reported previously in dendritic cells moving under agarose as they push upward against the gel (Gaertner et al., 2022). Unconfined macrophages in the same dish often demonstrated dorsal actin ruffles rather than actin puncta (**Fig. S1B**). Initial tests were conducted in setups with both well distances, in the absence of applied directional cues and with untreated glass (**Fig. 1B**, schematic at bottom). A population of cells is defined as migrating directionally if the mean FMIx ± 95% confidence interval does not encompass zero. To our surprise, macrophages migrating under the agarose in the close condition in Zone 2 had FMIx values that indicated persistent radial migration. The directional response of cells under agarose appeared to be due simply to entering a confined setting, regardless of their (+) or (-) orientation relative to the center media well (**Fig. 1C, Movies S1 and S2**). In this experimental setup there was no detectable difference between velocity, persistence or Euclidean distance between groups (**Fig. 1C, Fig. S2A**). These data suggest that macrophages move in a directionally persistent fashion upon entering a confined environment.

**Figure 1.**
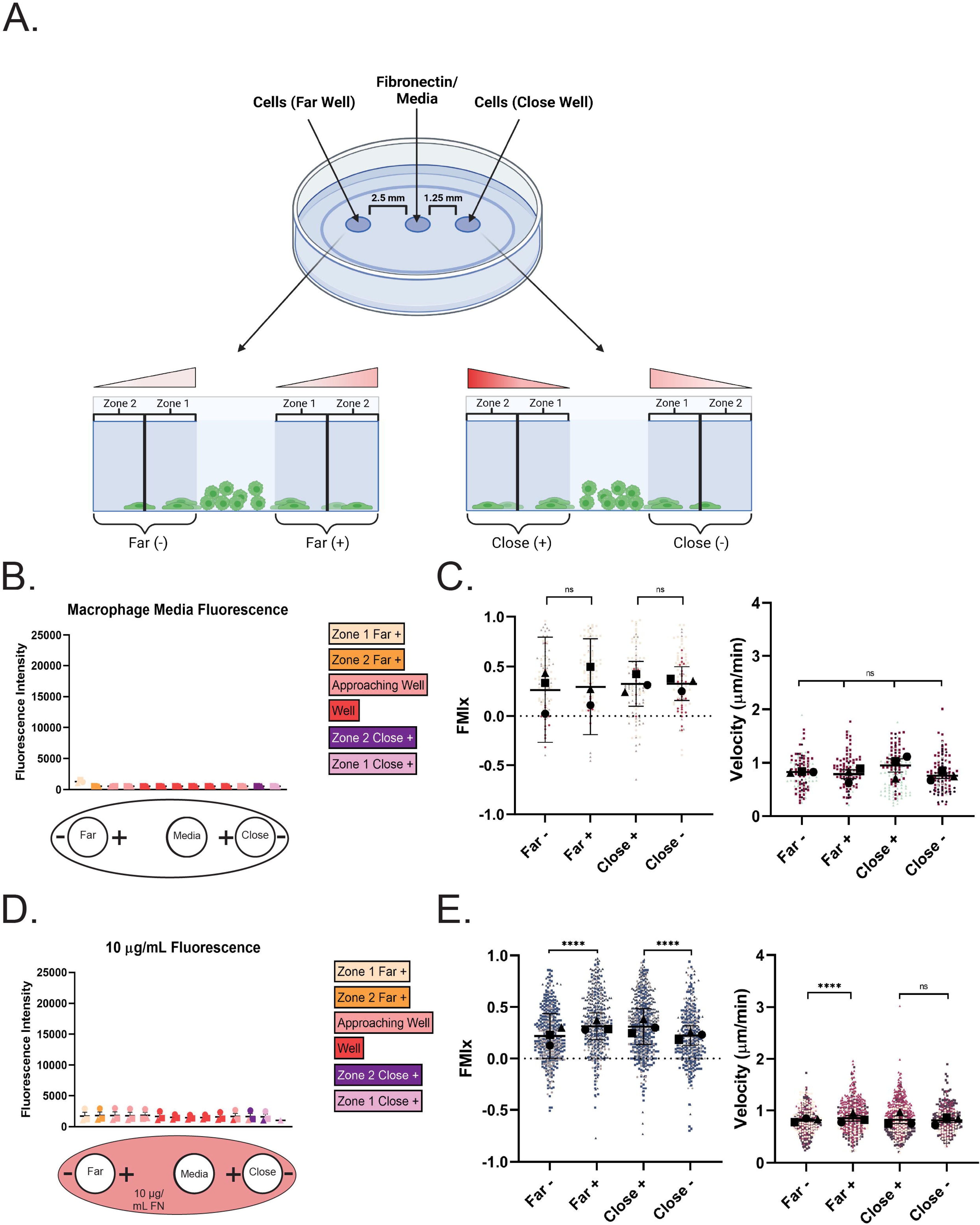
Transitioning to confinement induces directional migration in macrophages. (A) *Top*: Schematic of under-agarose model containing a ‘source’ well and cell wells on either side, generated via biopsy punch at discrete positions from the source well. *Bottom*: Cells migrating under the agarose move through Zones 1 and 2. The schematic depicts the direction and intensity of the haptotactic gradient above the areas of interest. The + and – indicate if the cells are moving towards or away from the center source well. This schematic was created with BioRender.com. (B) Macrophage migration under confinement with no ECM pre-coating. *Top:* Quantification of fluorescence across Zone 1 and Zone 2 at the end of the experiment (16 hours). Each point is the average fluorescence intensity (average grey) of each location in the chamber in N=3 experiments plotted with the SEM. *Bottom:* Diagram of experimental set-up with no ECM coating. Cells migrate either towards (+) or away (-) from the center well. (C) Migration relative to the center well is represented by FMIx (left) and velocity (right). Means and 95% confidence interval (for FMIx) or means and SEM (velocity) are represented with black symbols, and all cell migration tracks are plotted and each experimental run is color- and shape-coded (circle, square, or triangle). Statistical analysis was assessed with Kruskal–Wallis and Dunn multiple comparisons test. ns= not significant. Far-*n* = 123 tracks, Far+ *n* = 104 tracks, Close+ *n*= 139 tracks, Close-*n =* 123 tracks. These data were pooled from 3 independent experiments. (D) Macrophage migration under confinement with uniform pre-coating of 10 µg/mL FN. *Top*: Fluorescence quantification of dish pre-coated with 10 µg/mL RRX-FN. Quantification is performed the same way as stated in (B) at the 16-hour timepoint and represents the mean and SEM of 3 combined experiments. *Bottom*: Diagram of experimental set-up in which dishes were pre-coated with uniform 10 µg/mL FN. Media was again added to the center well, and macrophages were loaded into either the far or close well. Cells migrate either towards (+) or away (-) from the center well. (E) Migration relative to the center well is represented by FMIx (left) and velocity (right). Means and 95% confidence interval (for FMIx) or means and SEM (velocity) are represented with black symbols, and all cell migration tracks are plotted and each experimental run is color- and shape-coded (circle, square, or triangle). Statistical analysis was done by Kruskal–Wallis with Dunn multiple comparisons test. \*\*\*\**p* < 0.0001, ns = not significant. Far-*n* = 501 tracks, Far+ *n* = 460 tracks, Close+ *n*= 609 tracks, Close-*n =* 378 tracks. These data were pooled from 3 independent experiments.

Next, we were curious to know whether extracellular matrix alters confinement-induced directional migration. Uniform coatings of 10 µg/mL FN at this concentration are known to elicit random macrophage migration on 2D surfaces (Stinson et al., 2024). The fluorescence intensity across the glass surface was quantified, confirming that RRX signal was higher in these experiments than in the uncoated glass experiments (compare **Fig. 1B to Fig. 1D**). Once again, macrophages moved radially and with high directional persistence through Zone 2, hours after moving into a confined setting, indicated by the 95% confidence intervals that do not encompass zero (**Fig. 1E, Movies S3 and S4**). The directional response in the presence of FN appeared more robust than on uncoated glass, and macrophages migrating under agarose moved more persistently and faster than we have seen on the same concentration of FN in unconfined 2D culture (Stinson et al., 2024), where cells have an average speed of 0.4 µm/min and average persistence values around 0.3. It is worth noting that slight but statistically significant changes in mean FMIx values were noted, depending on which direction (+ or -) macrophages moved under the gel (**Fig. 1E**). Once again, cells entering confinement in the close well spacing demonstrated identical velocity, persistence and Euclidean distance values regardless of which side of the well they entered (+ or -, **Fig. 1E, Fig. S2B**). Unlike cells migrating under agarose on uncoated glass, cells entering confinement in the far well spacing on 10 µg/mL RRX-FN demonstrated slight but statistically significant differences in all measured values that depended upon whether they entered confinement from the + or – orientation (**Fig. 1E, Fig. S2B**). Another cue in the assay such as hydrostatic pressure, viscosity, or a weak media gradient diffusing from the central well might synergize with FN to slightly influence cellular behavior in the ‘far’ well spacing such that cells facing the center well (‘+’ condition) are slightly more directionally biased. To directly test the idea that adhesion prior to confinement is critical for the directional response, we injected macrophages under the agarose, which prevented them from adhering to RRX-FN prior to confinement. Macrophages moved much more randomly under these conditions (**Fig. S3A, Movie S5**). The FMIx was fixed at zero, reflecting that cells injected under agarose all have a different directional frame of reference (**Fig. S3A**). Additionally, the relatively low persistence and velocity values of this population are more reflective of cells migrating randomly in unconfined 2D contexts and do not replicate the higher values induced when adherent macrophages transition to a confined environment (**Fig. S3A**, compare to **Fig. 1E** and **Fig. S2A**). It is possible that cell crowding could explain the directional response in our assay, as macrophages in the rear could be driving leader cells forward. We used a removable barrier system to test this possibility. Upon barrier removal, leading cells migrated with mean FMIs just above zero and modest persistence values, reflecting a much weaker directional response compared to the 10 µg/mL FN under agarose experiments in which cells moved with higher FMI and persistence values (**Fig. S3B**, compare to **Fig. 1E**). Finally, we confirmed that confinement enhances migratory persistence in an agarose-independent adhesion system that uses PDMS micropillars to induce a defined confinement height between two glass surfaces (Liu et al., 2015). In this experiment, confined and unconfined cells were compared side by side. As this population was plated relatively sparsely, (similarly to experiments in **Fig. S3A**) cells did not have the same directional frame of reference. We focused primarily on how induced confinement of single cells affected migratory persistence. As with under agarose confinement, cells became more directionally persistent after micropillar confinement compared to unconfined counterparts, although the velocity of each group was similar (**Fig. S3C**). These findings suggest that the transition from an adherent 2D environment to ECM-containing confinement elicits a durable directional migratory response, though cell crowding may play a minor role. We next wondered whether applying a gradient of FN would synergize with the effect of confinement to further enhance the directional response.

### Haptotactic fibronectin gradients direct macrophage migration during confinement

To confirm a gradient was present after diffusion outward from the source well, RRX-FN fluorescent intensity was measured at locations between the source, far and close wells over the course of a 16-hour experiment (see schematic in **Fig. 1A**). Overall, the plot profile revealed increasing RRX intensity as a function of how close the location was to the central well where RRX-FN was loaded, and that the gradient persisted over 16 hours (**Fig. 2A**). We used total internal reflection fluorescence (TIRF) microscopy to confirm that RRX-FN was deposited directly on the glass surface, and that RRX intensity was higher in close wells than at a similar position in far wells (**Fig. 2B**). These findings confirm that our fibronectin gradients are stable, linear and reproducible.

**Figure 2.**
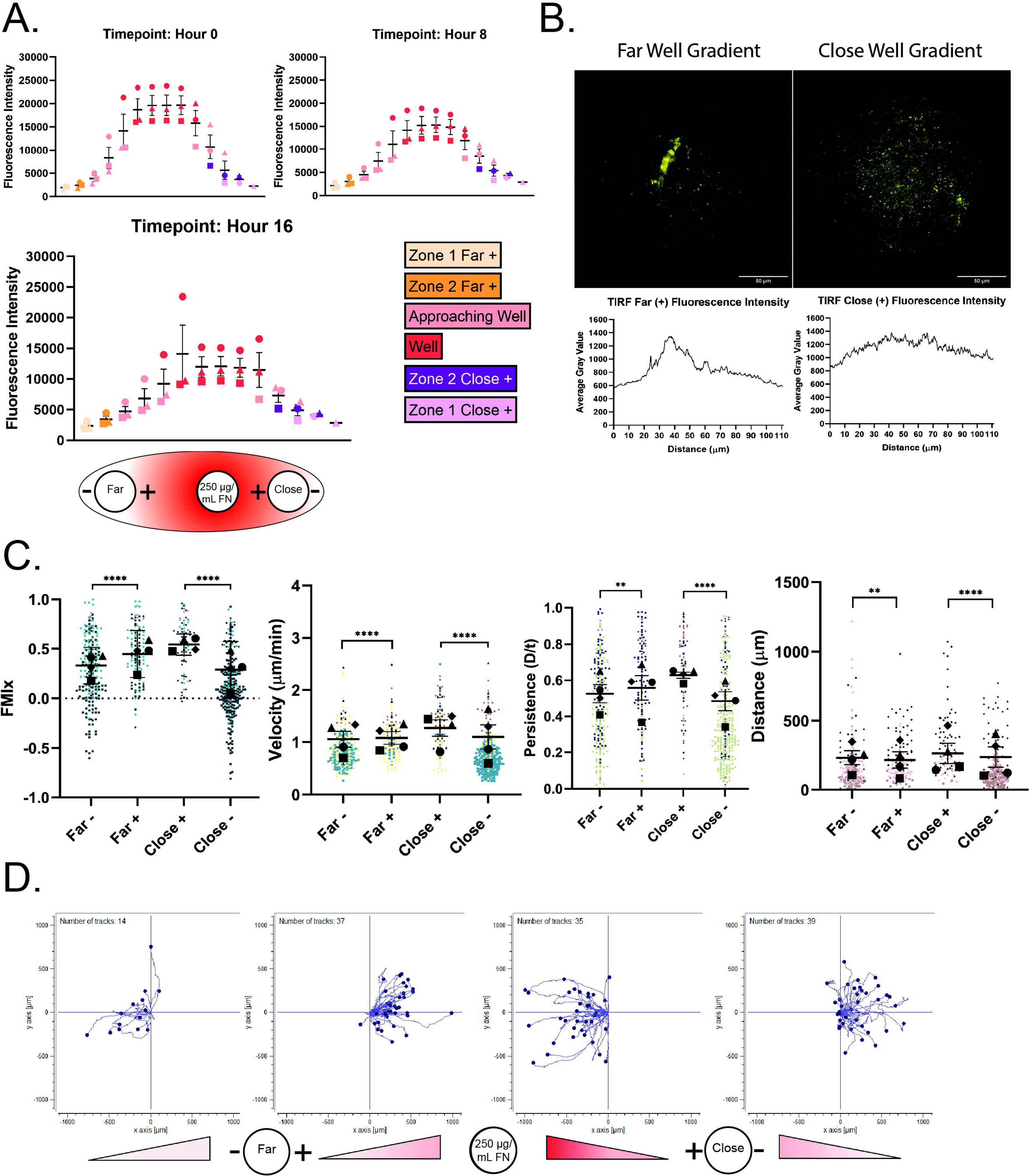
A haptotactic fibronectin gradient increases the macrophage directional response to confinement. (A) *Top:* Quantification of RRX-FN fluorescence gradient intensity over 16 hours. Colors correspond with the location in the well. Graph depicts the mean and SEM of N=3 experiments. *Bottom:* Diagram of experimental set-up depicts the gradient and source concentration of 250 µg/mL FN as it diffuses across the bottom surface of the dish. Media was again added to the center well and macrophages were loaded into either the far or close well. Cells migrate either towards (+) or away (-) from the center well. (B) Representative TIRF microscopy images of RRX-FN on the bottom of the dish (in Zone 1, just outside the far or close cell well) with corresponding line trace. The TIRF field of view is circular and is presented in its entirety in these images. Scale bar = 50 microns. (C) Migration relative to the center well is represented by (left to right) FMIx, velocity, persistence and Euclidean distance. Means and 95% confidence interval (for FMIx) or means and SEM (velocity, persistence, Euclidean distance) are represented with black symbols, and all cell migration tracks are plotted and each experimental run is color- and shape-coded (circle, square, diamond, or triangle). Statistical analysis was done by Kruskal–Wallis with Dunn multiple comparisons test. *\*\*\*\*p* < 0.0001, \*\**p* = 0.0090 (Persistence), \*\**p* = 0.0015 (Euclidean distance). Far-*n* = 224 tracks, Far+ *n* = 137 tracks, Close+ *n*= 87 tracks, Close-*n =* 276 tracks. These data were pooled from 4 independent experiments. (D) Representative migration track plots generated from the ibidi chemotaxis and migration tool FIJI plugin from a single imaging position. Each cell track is placed at the origin of the coordinate plane at time zero, with position tracked over a sixteen-hour time course. The labels on the bottom indicate the direction of the gradient relative to the distance and direction from the center well. The X and Y axis are in microns.

Presentation of a haptotactic gradient of FN (**Fig. 2A**, bottom) to cells entering confinement had a noticeable effect on macrophage migration characteristics. In this context, cells moving in the + and – orientations sense different parts of the gradient. Macrophages that enter confinement in the + direction are moving toward an increasing concentration of fibronectin. Macrophages that enter confinement in the – direction are moving against the haptotactic gradient. This is an important distinction, as this experiment can test whether haptotaxis and confinement synergize (the + condition) or whether confinement or haptotaxis is the ‘dominant’ cue (the - condition).

The transition to confinement still induces radial directional migration, as all populations have positive FMI values (**Fig. 2C**). However, both groups of cells moving along the gradient (+) had higher FMIx, velocity, persistence and Euclidean distance values than cells moving against the gradient (-) in both well spacings (**Fig. 2C, Movies S6 and S7**). It is also worth noting that cells moving against the gradient (-) in these experiments had more negative FMI values than cells moving with the gradient (+) (**Fig. 2C**, values below the dashed line at 0.0 FMI), suggesting that cells in the (-) condition might be turning around to follow the RRX-FN gradient. Individual tracks from one representative field of view (**Fig. 2D**) demonstrate qualitative differences between the (+) and (-) groups when a FN gradient is present. The tracks from the (+) condition tend to be straighter than those from the (-) orientation. Many cells moving against the gradient (-groups) moved more tangentially or even reverse direction (reflecting a negative FMI), indicating that they re-oriented to follow the FN gradient instead of continuing forward (**Fig. 2D**). Together, these data indicate that FN gradients are powerful directional cues that may synergize with confinement to heighten radial directional migration. These findings made us curious to know whether the transition into a confined environment could preserve a baseline of directional motility in a population of cells that cannot haptotax.

### Confinement induces a directional response in cells that cannot haptotax

Arp2/3-generated lamellipodial protrusions physically interact with haptotactic gradients and maintain directional persistence in this context (King et al., 2016; Rotty et al., 2017; Wu et al., 2012). In a related way, lamellipodia could help maintain the confinement-induced front/rear polarity required for directional persistence after cells move under agarose. We initially used agarose gels containing the Arp2/3 complex inhibitor CK-666 to interrogate this idea. CK-666 treatment did not impact confinement-induced directional migration as populations moving with (+) and against (-) the FN gradient (**Fig. S4A**, left) both moved with mean FMIx values well above zero (**Fig. 3A, 3B**). Unlike WT cells moving under agarose (**Fig. 2C**), the presence of a FN gradient did not boost the directionality of CK-666 treated cells moving with (+) the gradient compared to their counterparts moving against the gradient (**-**) (**Fig. 3B**). In fact, the (+) population movement was less directional and less persistent, and had a lower Euclidean distance than the (-) population (**Fig. 3B, Fig. S4A,** right), though migration velocity remained unchanged between the two groups (**Fig. 3B**). Non-muscle myosin II (NMII) has repeatedly been implicated in the response to confinement (Barbier et al., 2019; Liu et al., 2015; Lomakin et al., 2020), so we were curious whether inhibition of NMII by blebbistatin (bleb) would impair the directional response to confined haptotaxis. NMII inhibition did not affect macrophage haptotaxis, as FMI and persistence values were significantly higher in the ‘+’ direction compared to the ‘-‘ direction (**Fig. S4B**). This is in contrast to CK-666 treatment which affects both FMI and persistence of the ‘+’ population (**Fig. 3B**). Furthermore, velocity was unaffected by blebbistatin, reflecting previous findings on blebbistatin treatment of FN-adherent macrophages in 2D (Stinson et al., 2024) (**Fig. S4B**). These data suggest non-muscle myosin II is not required for macrophage directional persistence under confinement. We sought to probe the involvement of Arp2/3 complex more directly by comparing the motility of control and Arp2/3 knockout macrophages side by side as they migrated under agarose in response to a FN gradient.

**Figure 3.**
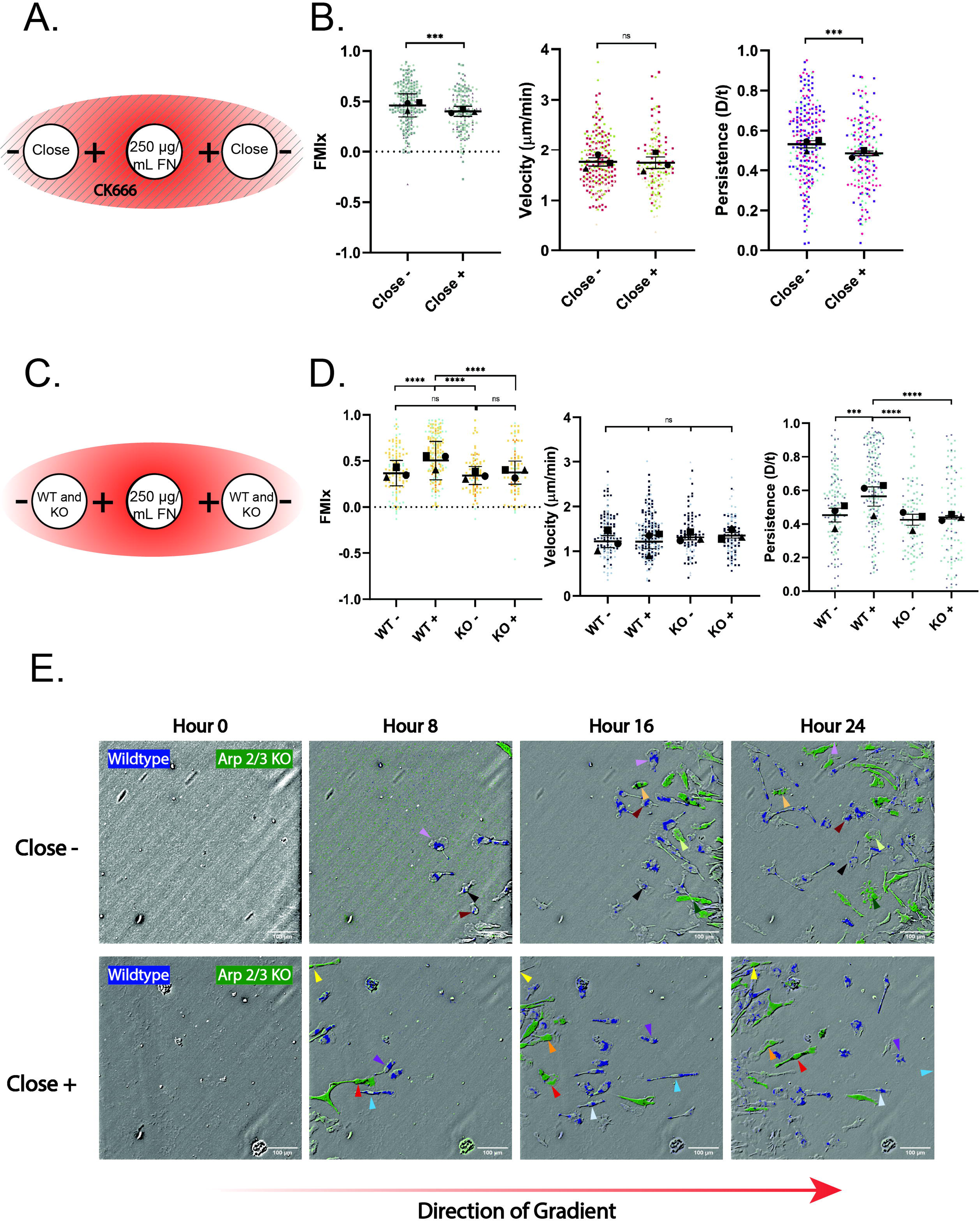
Arp2/3 complex disruption hinders haptotactic sensing but not confinement-induced directional migration. (A) Schematic depicting CK-666 experimental conditions. A 250 µg/mL FN gradient is generated as normal, but in this case the agarose is made up with 125 µM CK-666, and 60 µM CK-666 is included in center well, and both cell wells. Solid lines running through the schematic represent CK-666 polymerized into the gel. (B) Migration relative to the center well is represented by FMIx (left), velocity (middle), and persistence (right). Means and 95% confidence interval (for FMIx) or means and SEM (velocity, persistence) are represented with black symbols, and all cell migration tracks are plotted and each experimental run is color-and shape-coded (circle, square, or triangle). Statistical analysis was done by Mann-Whitney test. \*\*\**p* = 0.0002 (FMIx), \*\*\**p* = 0.0010 (Persistence), ns = not significant. Close-n = 250 tracks, Close+ n = 194 tracks. Data were pooled from 3 independent experiments. (C) Schematic depicting experimental conditions for WT vs KO (*Arpc2-/-*) macrophages migrating concurrently on 250 µg/mL FN gradient. (D) Migration of WT and KO (*Arpc2-/-*) macrophages toward the center well is represented by FMIx (left), velocity (middle), and persistence (right). Means and 95% confidence interval (for FMIx) or means and SEM (velocity, persistence) are represented with black symbols, and all cell migration tracks are plotted and each experimental run is color- and shape-coded (circle, square, or triangle). Statistical analysis was done by Kruskal–Wallis with Dunn multiple comparisons test. ****p < 0.0001, ***p < 0.0002, ns = not significant. WT-*n* = 128, WT+ *n* = 202, KO-*n* = 95, KO+ *n* = 97. These data were pooled from 3 separate experiments. (E) Representative images of differentially labeled WT (blue) vs KO (green) cell migrating against (close -) or along (close +) the fibronectin gradient. Representative KO (*Arpc2-/-*) cells migrating away from the gradient (close -) are marked with arrows colored light orange, light green, and dark green. KO (*Arpc2-/-*) cells migrating towards the gradient (close +) are marked with arrows colored red, orange, and yellow. Representative WT cells migrating away from the gradient (close-) are marked with arrows colored chestnut, brown, and light purple. WT cells migrating towards the gradient (close +) are marked with light blue, purple, and white. Scale bar = 100 microns. The direction of the FN gradient is schematically depicted below.

Tamoxifen-inducible loss of the Arpc2 subunit of the complex generates cells that completely lack Arp2/3 function (Paul et al., 2017; Rotty et al., 2017; Rotty et al., 2015), and therefore lack the ability to generate branched actin networks. WT and *Arpc2-/-* macrophages (**Fig. S5A**) were labeled with green (*Arpc2-/-*, or ‘KO’) or blue (WT) cell tracker dye to discriminate between them so that the two populations could be plated together in the central well (**Fig. 3C**). As expected, WT macrophages moving with the FN gradient (+) (**Fig. S5B**) had a significantly higher FMIx value compared to WT macrophages moving against the FN gradient (-), as well as compared to both *Arpc2-/-* populations (**Fig. 3D; Movies S8 and S9**). The other three populations, WT (-), *Arpc2-/-* (-) and *Arpc2-/-* (+), all had statistically similar FMIx values to one another (**Fig. 3D**), though with positive FMIx values of these populations that indicate the capacity for radial migration upon entering confinement. Notably, *Arpc2-/-* migration velocity remained similar to WT cells, regardless of orientation relative to the gradient (**Fig. 3D**), indicating that the altered FMIx was not due to loss of migratory capacity. The more meandering migration of *Arpc2-/-* (+) macrophages compared to WT (+) macrophages is also illustrated by the decreased persistence (**Fig. 3D**) and Euclidean distance (**Fig. S5C**) of the KO population. **Figure 3E** shows two example populations labeled with green (*Arpc2-/-*) or blue (WT) tracker dye. Arrows denote individual cells migrating across the time series. The top series demonstrates cells moving against (-) the fibronectin gradient, while the bottom series demonstrates cells moving with (+) the gradient. These images provide examples of WT macrophages moving radially in response to the FN gradient, compared to *Arpc2-/-* macrophages migrating beside them that meander more, because they did not sense the FN gradient (**Fig. 3E**, bottom). WT and *Arpc2-/-* macrophages migrating against (-) the gradient can also were seen to meander more (**Fig. 3E**, top). Though they lack lamellipodia, *Arpc2-/-* macrophages generated a combination of blebs and filopodial protrusions under agarose, similarly to how they generate protrusions in 2D environments (Rotty et al., 2017) (**Fig. S6A-B; Movies S10-11**). These compensatory protrusions appear to be adequate for macrophage translocation in confined environments, but are inadequate for haptotactic sensing. While both CK-666 and genetic deletion of Arp2/3 caused migration phenotypes consistent with loss of FN haptotaxis, cellular behavior in each case was subtly different. The most notable example is that CK-666 (+) cells had lower FMI and persistence values than CK-666 (-) cells, while the *Arpc2-/-* cells migrated with similar FMI and persistence values in both the (+) and (-) orientations. These differences may be accounted for by how these two disruptions act on Arp2/3 function. CK-666 acutely impairs Arp2/3, which renders them immediately incapable of generating branched actin networks. On the other hand, tamoxifen-inducible deletion of *Arpc2* occurs more gradually, thereby allowing these cells to compensate by upregulating other actin assembly pathways to try to maintain homeostasis (Burke et al., 2014; Rotty et al., 2015; Suarez et al., 2015). Acute versus long-term disruption of Arp2/3 complex has been previously shown to have subtle differences in phenotypic degree, while still yielding consistent overall insight (Rotty et al., 2017). Taken together, the experiments in the present study suggest that haptotactic FN gradients boost the directional effect of entering confinement. These data also reveal that the Arp2/3 complex is not required for confinement-induced radial migration. Therefore, lamellipodial protrusion may not be required to establish or maintain the front/rear polarity and directional response maintained after transitioning into confinement. However, our data strongly suggests that cells capable of simultaneously sensing multiple cues migrate more efficiently than counterparts responding to a single cue. This may have important implications *in vivo*, given the number of potential stimuli present in the microenvironment.

## DISCUSSION

The current study suggests that the transition to confinement is sufficient to induce a persistent directional migratory response in bone marrow-derived macrophages. It is easy to imagine that the transition to a confined environment may be a relevant migratory stimulus in several *in vivo* contexts, such as extravasation of circulating leukocytes and tumor cells into peripheral tissues. It has been known for many years that mechanical confinement spontaneously induces migratory behavior in cancer cells (Irimia and Toner, 2009), and increases their directional persistence (Mak et al., 2011). In addition, increasing confinement has been implicated in driving mesenchymal to amoeboid transitions (Holle et al., 2019), which seems to depend on both the microenvironment and specific cytoskeletal regulators (Gabbireddy et al., 2021). Leukocytes are similarly regulated by confinement. One striking example is the recent finding that Th1 T cells become migratory under confinement, and use integrin-independent adhesion to ECM to move (Caillier et al., 2024). Like some cancer cells, monocytic THP-1 cells become amoeboid under confinement and require myosin enrichment at the cell rear to move efficiently (Ullo et al., 2024). Mature, amoeboid dendritic cells also require non-muscle myosin II to efficiently migrate through dense, confining collagen gels and *in situ* ECM (Barbier et al., 2019). As confinement levels increase, myosin II activation increases in response to nuclear deformation, which allows immature dendritic cells and cancer cells to polarize and migrate more efficiently (Lomakin et al., 2020). Similar observations were concurrently made using a zebrafish model (Venturini et al., 2020). However, several studies also demonstrate the capacity of some cells to migrate in a myosin-independent fashion under confinement (Balzer et al., 2012; Liu et al., 2015; Stroka et al., 2014). The data we present here indicates that macrophages under agarose do not require myosin II to move radially in response to confinement, or to haptotax. However, additional myosin II-focused studies will need to be conducted on macrophages, as there may be a context-dependent role for myosin II under confinement.

Confinement-induced directionality appears to be much stronger if FN is present. This implies that adhesion-dependent signaling is important for this response. While microchannel confinement is known to suppress focal adhesions in cancer cells (Balzer et al., 2012; Holle et al., 2019), adhesion signaling can be a critical component of the cellular response to confinement (Hung et al., 2013; Reversat et al., 2020). In addition, the process of integrin-dependent haptokinesis has recently been identified as an important factor for macrophage exploration of 3D environments containing efferocytosis cues (Paterson and Lammermann, 2022). Though there are likely physical differences between under agarose migration and 3D migration through dense ECM, a similar system could be at play when cells transition into an adhesive, confining environment. However, leukocytes are capable of migrating efficiently through the interstitial space even with compromised integrin-mediated adhesion (Lammermann et al., 2008) and can overcome loss of adhesion by using topological cues for guidance (Reversat et al., 2020). Therefore, it is possible that other strategies like polarized ion and water channels creating an ‘osmotic engine’ (Stroka et al., 2014), or microtubule leading edge polarization (Balzer et al., 2012) may also contribute.

Haptotactic FN gradients increased macrophage FMI, persistence, cell speed, and Euclidean distance after cells move under agarose, compared to cells in the same experiment that moved against the gradient. These data suggest that haptotaxis synergizes with confinement to induce a more robust migratory response. Haptotaxis requires stable leading edge lamellipodia that physically sense substrate-bound cues (King et al., 2016; Wu et al., 2012). The Arp2/3 complex generates the branched actin network that induces lamellipodial protrusion, and Arp2/3 is a major effector of haptotaxis in both 2D and 3D (King et al., 2016; Rotty et al., 2017; Wu et al., 2012). Similarly, Arp2/3 disruption renders macrophages unresponsive to haptotactic gradients under agarose, as they migrate less persistently and directionally than WT cells plated side by side on the same gradient. Nonetheless, the *Arpc2-/-* macrophages retain a positive FMI value, suggesting that they still move directionally in response to entering confinement. These data imply that the haptotactic and confinement responses are separable, and may be driven by synergistic signaling and cytoskeletal regulation when cells respond simultaneously to both cues.

It is likely that many cell types integrate multiple cues as they carry out their physiological functions. This may be relevant for many potential reasons. First, integrating distinct cues could lead to more effective directional responses *in vivo*. There is experimental support for this idea. For example, mature dendritic cells migrate persistently in 3D collagen gels along carbon microfibers coated with CCL21 and ICAM-1 in the presence of soluble CCL21 or CCL19 gradients, even if this meant forsaking a direct path along the chemokine gradient (Schumann et al., 2010). Such an ability to use topological and/or haptokinetic cues to continue migrating could allow cells to efficiently move around physical barriers like hair follicles in the skin. Related to this, the ability to integrate multiple cues may keep cells moving toward their destination if they lose the ability to sense a single cue, rather than backtracking or pausing. Our experimental results are consistent with this idea, as haptotaxis-deficient *Arpc2-/-* macrophages retain the ability to move with a positive FMI upon entering confinement, though less directionally than their WT counterparts that haptotax. In addition, different cues may be available to the ‘first responders’ to a site of action *in vivo* than are available to later waves of recruited cells, though the later arriving cells are likely just as important as the first wave. The first responders also likely alter the microenvironment as they move through it, depositing or removing cues, or physically altering the matrix available to trailing cells. Thus, it is advantageous for cells to be able to sense and respond coherently to a complex and shifting microenvironment. Finally, this adaptability may be required for cells to accomplish distinct phases of their *in vivo* function. Compelling examples come from recent work on dendritic cells. Before they mature, unstimulated dendritic cells (DCs) go through periods of migration and halting that are characterized by actomyosin enrichment at the rear during migration and the front during halting (Chabaud et al., 2015). Actomyosin enrichment at the front during this period corresponded to macropinocytic sampling, similar to how DCs scan their microenvironment for antigen (Chabaud et al., 2015). At the same time, these immature DCs are resistant to barotactic hydraulic cues due to their macropinocytic ability (Moreau et al., 2019). On the other hand, DC maturation via TLR stimulation makes them responsive to chemotactic cues, which corresponds with suppression of macropinocytosis due to localization of actomyosin at the rear of migrating cells (Chabaud et al., 2015; Vargas et al., 2016). Likewise, mature DCs become barotactic (Moreau et al., 2019). The shift toward migration upon maturation makes sense, as stimulated DCs need to find their way into lymph nodes rather than continue sampling their immediate environment for antigen. Indeed, haptotactic gradients of CCL21 have been noted *in vivo* and play a major role in recruiting mature DCs to lymph vessels (Weber et al., 2013). It is clear from these and other examples that the ability of cells to integrate distinct cues from their microenvironment is likely managed in a context- and tissue-specific fashion.

In summary, bone marrow-derived macrophages move in a highly directional fashion upon entering a confined environment, and inclusion of a FN gradient increases the directional response further. It is possible that the transition to confinement induces actomyosin polarization at the cell rear, similar to mature DCs (Vargas et al., 2016) or to neutrophils migrating directionally *in vivo* in response to laser wounds (Georgantzoglou et al., 2022) allowing for persistent directional migration to occur, though this is difficult to reconcile with the blebbistatin experiments in the present work. Extravasation into a dense tissue from the circulation may similarly induce an inherent directional response, thereby allowing cells to move persistently into a target tissue without needing to carefully integrate multiple microenvironmental cues. Once in the target tissue, cells could then respond to microenvironmental cues to further tune their migration, differentiation or proliferative behaviors depending on how they integrate localized cues. It will be interesting to determine the molecular drivers of confinement-induced migratory persistence to more fully understand how this process interacts with other directional cues and to interrogate this process in physiological and pathological responses *in vivo*.

## MATERIALS AND METHODS

### Bone Marrow Derived Macrophage Culture

BMDMs used in this study come from a mixed strain (though predominantly C57BL/6) of mice harboring a conditional Arpc2 allele. This allele consists of LoxP sites flanking exon 8 of the gene encoding the p34/Arpc2 subunit of the Arp2/3 complex (Rotty et al., 2017; Rotty et al., 2015). In addition, these cells harbor Ink4a-/-; Arf-/-alleles, which allow them to persist in culture without becoming senescent. This is a defined genetic event rather than a spontaneously arising oncogenic immortalization. Finally, these cells also harbor a Rosa26-CreER transgenic allele that allows for conditional deletion of Arpc2 exon 8 in the presence of tamoxifen. Macrophages were cultured in complete cell culture media (referred to as macrophage media from here on) composed of 70% DMEM complete (Thermo, 11054-020) + 10% Fetal Bovine Serum (Sigma, 12306C-500ML) + 1% Glutamax (Thermo, 35050061) and 30% L929-conditioned media (for a source of M-CSF). Macrophages were maintained in macrophage media at 37°C, 90% humidity, and 5% CO2. For passaging at 60-90% confluency, cells were incubated at 4°C for 10 minutes with pre-chilled 0.5 mM EDTA (Thermo, AM9260G). At the end of the incubation, EDTA was aspirated and macrophages were scraped directly into macrophage media and re-plated for passaging and/or experimentation. Cells not treated with tamoxifen are functionally wild-type for the duration of their time in culture. To generate Arp2/3 knock out cells, cells were treated with 2 µM 4-hydroxytamoxifen (Sigma, H7904-5MG) in macrophage media for 5 days. The generation of genetic Arpc2 knock outs in this line of immortalized bone marrow derived macrophages has been previously published (Ronzier et al., 2022; Rotty et al., 2017). Arpc2 (Millipore Sigma, 07-227-I-25UG) and GAPDH (Thermo, AM4300) antibodies were commercially sourced. Cells are tested for mycoplasma using a commercial testing kit (Invivogen, rep-mysnc-50).

### Agarose Plate Preparation

5 mL of 2x Hank’s Balanced Salt Solution (Sigma, H1387-10X1L) and 10 mL of macrophage media were combined and placed in a 68 °C bath for 1 hour. 0.24 g of low gelling temperature agarose (Sigma, A9045-5G) was added to 5 mL of sterile water (Corning, 25-055-CV) and heated in a microwave until fully dissolved. The two solutions were combined to create a warm 1% agarose mixture. When present, cascade blue-labeled 3000 M.W. dextran (Thermo Fisher, D7132) was added to the agarose prior to pouring. 3 mL of this mixture was poured into a 35 mm glass bottom dish containing a 20 mm micro-well (Cellvis, D35-20-1.5-N) and allowed to solidify for 90 minutes at room temperature. The plates were not pre-coated with substrate, except in the case of the 10 µg/mL uniform fibronectin experiment, which involved a 1-hour incubation of the Rhodamine RRX-fibronectin (Cytoskeleton, FNR01-A) in the micro-well at 37 °C prior to agarose addition. Plates were wrapped in parafilm and stored at 4 °C for up to one month, after which point the gel’s integrity could be compromised.

### Template Dish Preparation

A 3.5 mm biopsy punch tool (Robbins Instruments, RBP-35) was used to generate wells in the agarose gel. Punches were created and then excess gel was aspirated to create each well. In order to create chambers with the same distance between the well containing the haptotactic cue or control media, a reusable template was created using another 35 mm glass bottom dish. For experiments containing a close and far condition, the bottom of a 35 mm glass bottom dish was marked with an ethanol-resistant marker to indicate the size of the source well hole-punch. Then, the close and far wells were marked so that the edges of the source well and the cell-containing wells were 1.25 and 2.5 mm away, respectively. The agarose-coated dish to be used in a migration assay was placed on top of the template dish, so that the circles marking well location were visible through the bottom of the dish. For experiments with two close wells, a second template was created with only 1.25 mm well spacing.

### Cell Preparation

Macrophages were labeled with CellTracker Green CMFDA Dye (Invitrogen, #C7025) using a 1:4000 dilution in 1x Cell Culture Phosphate Buffered Saline (hereafter referred to as PBS) (Corning, MT21040CV) for 5 minutes at room temperature. Lifeact-mScarlet was stably transduced into macrophages using the previously described LentiLox system (Rotty et al., 2017), followed by FACS-based selection for red fluorescence. After labeling or transduction, the cells were treated with 0.5mM EDTA (Invitrogen, AM9260G) and scraped into 1 mL of macrophage media. Once counted, 100,000 cells were transferred to 2 separate eppendorf tubes (200,000 cells total) and spun down at 1000xg for 3.5 minutes. Excess media was aspirated, and cells were resuspended in 20 µL of macrophage media. These suspensions were then transferred to each of the close and far wells in the prepared agarose dish.

### Time-lapse Migration Video Capture and Analysis

<u>Time-lapse video microscopy</u> was conducted on the Olympus IX83, and an INU incubation system controller from Tokai Hit was used to maintain cells in a stable environment at 37°C, 90% humidity, and 5% CO2. Under-agarose chambers were placed into the environmental chamber. A 20X air objective was employed during overnight time lapse imaging. Cells were then imaged via relief contrast and the appropriate fluorescent channel for the chosen tracker dye. A total of 32 positions were selected to assay macrophage haptotaxis for each experiment: 4 non-overlapping sites from the top to the bottom of the edge of a cell well (Close or Far), which composed Zone 1, 4 more non-overlapping sites at the same Y position but shifted approximately 665 μm (1 field of view) away, which composed Zone 2. This process was repeated on the other side of the same well, and then on both sides of the second cell well. Images were captured at 10-minute intervals for 16 hours for the 250 μg/mL haptotaxis assay, macrophage media assay, and 10 μg/mL assay. The CK666 and Arp2/3 KO assays were imaged for 24 hours. Fiji (ImageJ) was used to calculate macrophage FMIx, velocity, Euclidean distance, and to generate migration plots. Cell tracks were generated with a combination of the TrackMate plugin for automated tracking and the Manual Tracking plugin. Cell tracks were then imported into the Ibidi Chemotaxis and Migration Tool plugin to quantify migration characteristics. FMIx is calculated by dividing the difference between the starting and ending X coordinate of a cell track value by the accumulated distance. Euclidean distance is the straight-line length between the first and last X,Y coordinate of a cell track. Persistence is the ratio between a cell’s Euclidean distance (D) and its total path length (t), with values closer to 1 indicating more directional persistence. Note that persistence is not the same as FMIx as the persistence values do not consider any external frame of reference (e.g. an external gradient). *Arpc2-/-* macrophages were also subjected to time-lapse imaging at 40x using a DIC objective. In these experiments, cells were imaged for 10 minutes, with 10 second intervals. When necessary, XY drift was corrected in FIJI with the 3D drift correction plugin under the ‘Registration’ plugin heading. Cells were then scored for filopodia, blebs, and large leading edge blebs by eye. The average fraction of each protrusion type as a share of the total number of protrusions was then reported. <u>Confocal microscopy</u>: A Nikon inverted spinning disk confocal microscope outfitted with an EMCCD camera (Yokogawa) and 60x Plan Apo 1.4 NA oil objective was used to image the agarose-dextran confinement of Lifeact-Scarlet expressing macrophages. NIS-Elements software controlled the microscope during image acquisition. Z-sections were taken at 0.3 micron intervals, and side projections were constructed using the NIS-Elements software.

### Rhodamine RedX (RRX) Fibronectin (FN)/Media Quantification

RRX-FN (Cytoskeleton, FNR01-A) or macrophage media was allowed to spread under agarose from the central chamber for 1 hour at 37 °C prior to each migration assay. 12-14 images were taken across the length of the chamber between the (+) regions of the cell wells during migration runs. Fluorescence power was set to 20% and exposure time was set to 300 ms. Each position was opened in ImageJ, and the DsRed channel was isolated, and Brightness/Contrast was set manually to a minimum value of 0 and a maximum value of 20000. Using Edit -> Selection -> Specify, a 500W x 1H pixel line is created at the X coordinate 262 and the Y coordinate 512 to measure the fluorescence intensity across the center of the image. A Gray Value plot was generated, and the data was saved to Excel and then loaded into Graphpad Prism.

### Arp2/3 KO (Arpc2-/-) vs WT Macrophage Under-Agarose Haptotaxis Assay

Agarose chambers were prepared as previously described. During RRX-FN diffusion, *Arpc2-/-* and WT macrophages were differentially labeled with Cellbrite Cytoplasmic Membrane Dye Blue (WT) (Biotium, #30024) or CellTracker Green CMFDA Dye (*Arpc2-/-*) (Invitrogen, #C7025), as previously described. The blue dye was observed to be secreted at a low level by WT cells and taken up by green *Arpc2-/-* cells at a low rate, yielding some doubly labeled cells. Conversely, we saw no evidence of green dye secretion or transfer. Thus, green cells as well as green cells with low level blue staining were both identified as *Arpc2-/-* while cells labeled with only the blue dye were identified as WT.

### CK-666 and Blebbistatin Under-Agarose Haptotaxis Assay

The small molecule Arp2/3 complex inhibitor CK-666 (Abcam, ab141231) and blebbistatin (S-nitro blebbistatin; Fisher Scientific, NC0664123) were polymerized with agarose at a final concentration of 125 µM CK-666 or 30 µM Blebbistatin. The agarose chamber was then prepared as previously described with one source well and two close wells, and a RRX-FN established as previously described. Cells were resuspended in 20 µL of macrophage media also containing 60 µM CK-666 or 15 µM Blebbistatin before being added to each of the close wells. 60 µM CK-666 or 15 µM Blebbistatin in macrophage media was also present in the source well.

### Removable chamber migration assays

Glass bottomed dishes were treated with 10 µg/mL FN as previously described. At the same time, silicone inserts containing 0.4 mm wells (ibidi, 80486) were removed from the glass dish they were shipped on and placed in a sterile secondary container. The same punch biopsy device used to make agarose gel wells was used to punch four holes around the existing 0.4 mm wells. This modified insert was then placed back on top of the dry FN-plated glass and pressed down gently to make a liquid-tight seal with the fibronectin-coated glass. Macrophages were prepared as previously described. Cell pellets were resuspended in 80 µL of macrophage media. 20 µL of this suspension was added to each of the four wells produced by the biopsy punch. After aspirating excess media, the insert was carefully lifted from the dish with forceps. Adherent cells were washed, and 2 mL macrophage media was added to the dish prior to the start of time lapse imaging.

### Static confinement device

Macrophages were plated on 10 µg/mL FN in a glass-bottomed dish, as previously described, and placed in a tissue culture incubator at 37 °C overnight to allow cells to adhere to the substrate. The following morning, the dish was placed into an adjustable one well static confiner (4Dcell, CSOW110), which was set using confinement coverslips containing PDMS micropillars (4Dcell, CONFCS 110-12) that maintained a 5-micron confinement height. Cells were imaged immediately after the initiation of confinement, in time lapse experiments similar to those outlined above. Cells under the confinement coverslip (confined) were compared to cells at the edge of the glass surface that were outside the radius of the confinement coverslip and were therefore unconfined.

### Statistical Analysis

The Kruskal–Wallis with Dunn multiple comparisons test was used to assess significance in experiments where a normal distribution of the dataset could not be assumed. When only two experimental conditions were tested, we used Mann–Whitney tests when we could not assume normality. Unpaired t tests and ANOVAs were used when normality tests indicated a normal distribution of the data. All statistics were calculated using GraphPad Prism, and significance was assumed if p ≤ 0.05. More information on each statistical test can be found in the relevant figure legend panel.

## LIST OF ACCOMPANYING SUPPLEMENTARY MATERIALS

1) Supplemental movie 1: Macrophages in Zone 2 (+) migrating with media only.

2) Supplemental movie 2: Macrophages in Zone 2 (-) migrating with media only.

3) Supplemental movie 3: Macrophages in Zone 2 (+) migrating on 10 ug/mL uniform fibronectin.

4) Supplemental movie 4: Macrophages in Zone 2 (-) migrating on 10 ug/mL uniform fibronectin.

5) Supplemental movie 5: Macrophages in Zone 2 migrating on 10 ug/mL uniform fibronectin, after being directly injected under agarose.

6) Supplemental movie 6: Macrophages in Zone 2 (+) migrating on 250 μg/mL haptotactic gradient.

7) Supplemental movie 7: Macrophages in Zone 2 (-) migrating on 250 μg/mL haptotactic gradient.

8) Supplemental movie 8: Mixed population of WT and KO macrophages in Zone 2 (+) migrating on 250 μg/mL haptotactic gradient.

9) Supplemental movie 9: Mixed population of WT and KO macrophages in Zone 2 (-) migrating on 250 μg/mL haptotactic gradient.

10) Supplemental movie 10: Example of bleb-producing *Arpc2-/-* macrophage migrating under agarose on 250 μg/mL haptotactic gradient, 40x magnification with 10s intervals.

11) Supplemental movie 11: Example of filopodia-producing *Arpc2-/-* macrophage migrating under agarose on 250 μg/mL haptotactic gradient, 40x magnification with 10s intervals.

## ACKNOWLEDGEMENTS

We thank the members of the Rotty Lab for helpful discussions during research development, and to Dr. Prasanna Satpute-Krishnan for assistance with live cell confocal imaging. This project is sponsored by the Uniformed Services University of the Health Sciences (USU); however, the information or content and conclusions do not necessarily represent the official position or policy of, nor should any official endorsement be inferred on the part of, USU, the Department of Defense, or the U.S. Government.

## AUTHOR CONTRIBUTIONS

M.S.: experiments and planning, data analysis, writing and editing. S.P.: experiments and planning, data analysis, writing and editing. E.C.: experiments, data analysis, editing. J.R.: Project oversight, experiments and planning, writing, editing, funding. All authors had the opportunity to review and comment on the manuscript prior to submission.

### DATA AVAILABILITY STATEMENT

All primary data will be openly available upon request.

---

**Supplemental Figure 1.**
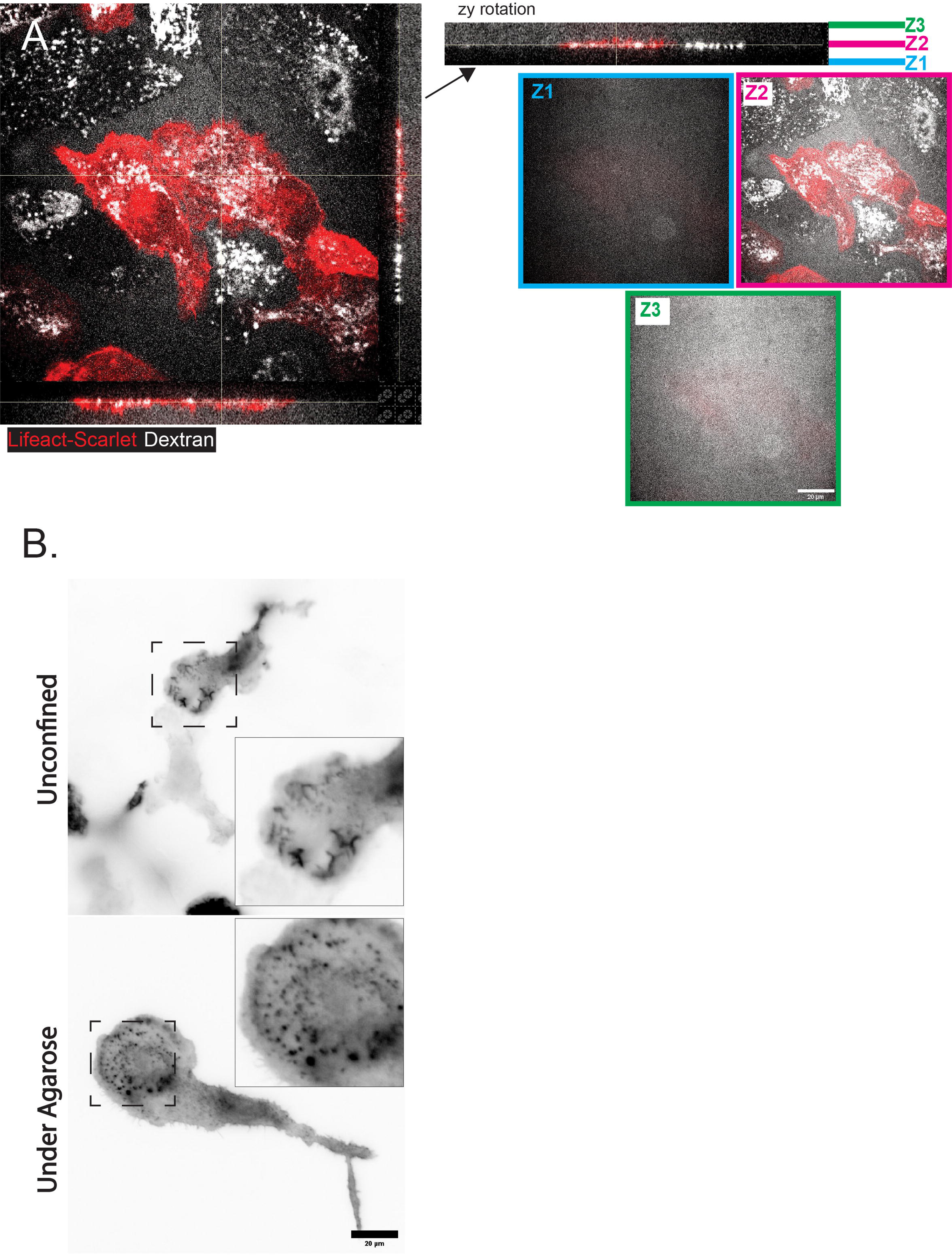
Additional quantification related to Figure 1 part 1. (A) *Left*: Confocal image of Lifeact-Scarlet expressing cells (red staining) within an agarose gel containing fluorescent dextran (white staining). Side projections corresponding to the vertical and horizontal lines across this image are at the right and bottom of this image, respectively. *Right*: The vertical side projection has been rotated (top) and color-coded markers have been included to mark the bottom (Z1), middle (Z2) and top (Z3) of the image series consisting of 43 total images, which is composed of z slices taken at 0.3-micron intervals. Of these, 17 slices span the width of the indicated cell. The images corresponding to the three representative positions (Z1, Z2, Z3) are taken from the 43 total image slices and are reproduced with color-coded borders. As with the image on the left, dextran is colored white and Lifeact-Scarlet is colored red. Scale Bar = 20 microns. (B) Example images of LA-Scarlet staining in live macrophages in unconfined (top) or agarose-confined (bottom) settings. Scale bar = 20 microns and both images are at the same scale. Insets have been maintained at the same scale relative to each other.

**Supplemental Figure 2.**
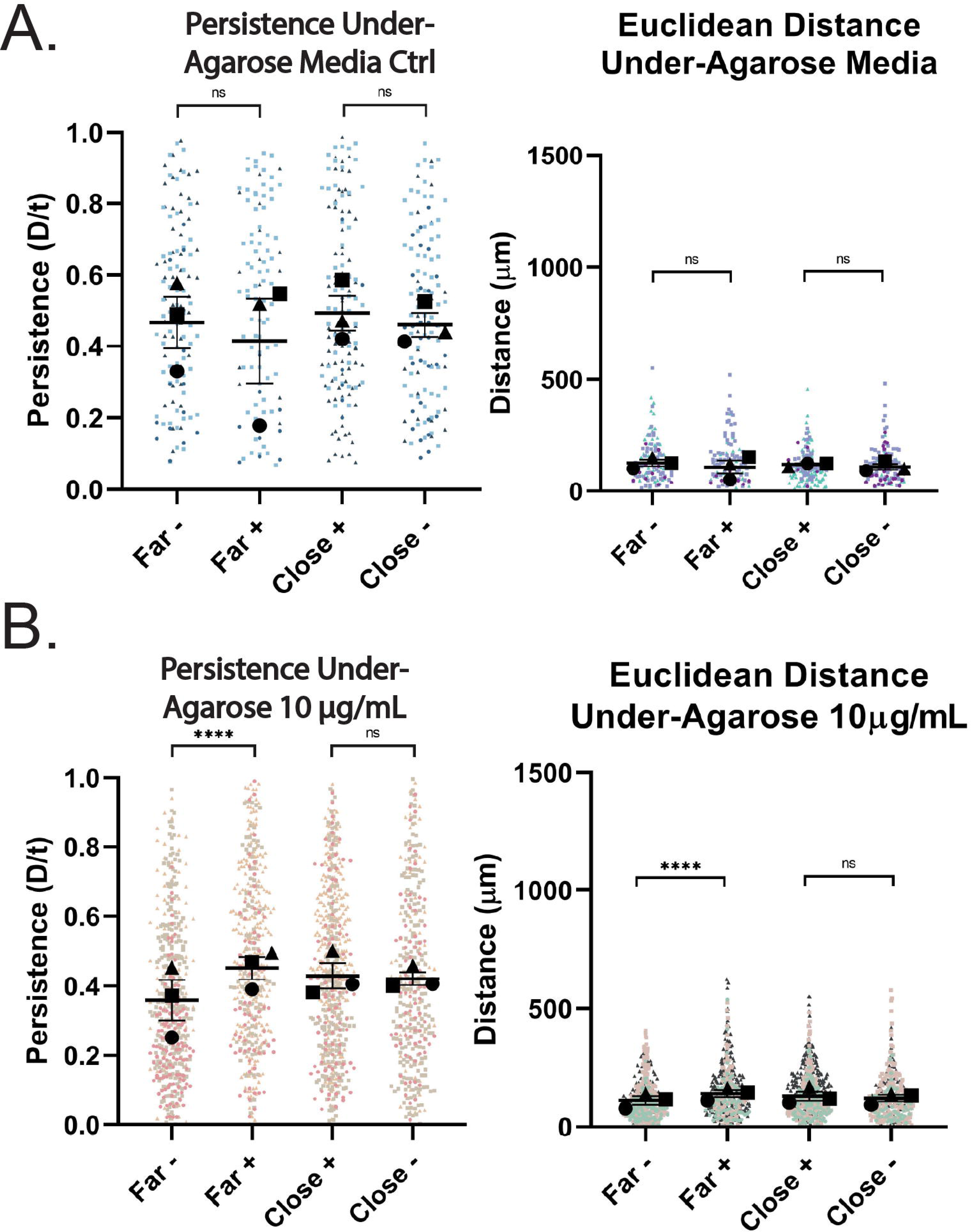
Additional quantification related to Figure 1, part 2. (A) Persistence (left) and Euclidean distance (right) of Far +/- and Close +/- macrophages migrating under agarose with media only. Both values were calculated from the same dataset as **Fig. 1C**. Experimental means and SEM are represented with black symbols and all cell migration tracks are plotted and each experimental run is color- and shape-coded (circle, square, or triangle). Statistical analysis was assessed with Kruskal–Wallis and Dunn multiple comparisons test. Ns= not significant. Far-*n* = 123 tracks, Far+ *n* = 104 tracks, Close+ *n* = 139 tracks, Close-*n =* 123 tracks. These data were pooled from 3 independent experiments. ns = not significant (B) Persistence (left) and Euclidean distance (right) of Far +/- and Close +/- macrophages migrating under agarose in the presence of uniform 10 µg/mL RRX-FN. Both values were calculated from the same dataset as **Fig. 1E**. Experimental means and SEM are represented with black symbols and all cell migration tracks are plotted and each experimental run is color- and shape-coded (circle, square, or triangle). Statistical analysis was assessed with Kruskal–Wallis and Dunn multiple comparisons test. \*\*\*\**p* < 0.0001, ns = not significant. Far-*n* = 501 tracks, Far+ *n* = 460 tracks, Close+ *n*= 609 tracks, Close-*n =* 378 tracks. These data were pooled from 3 independent experiments.

**Supplemental Figure 3.**
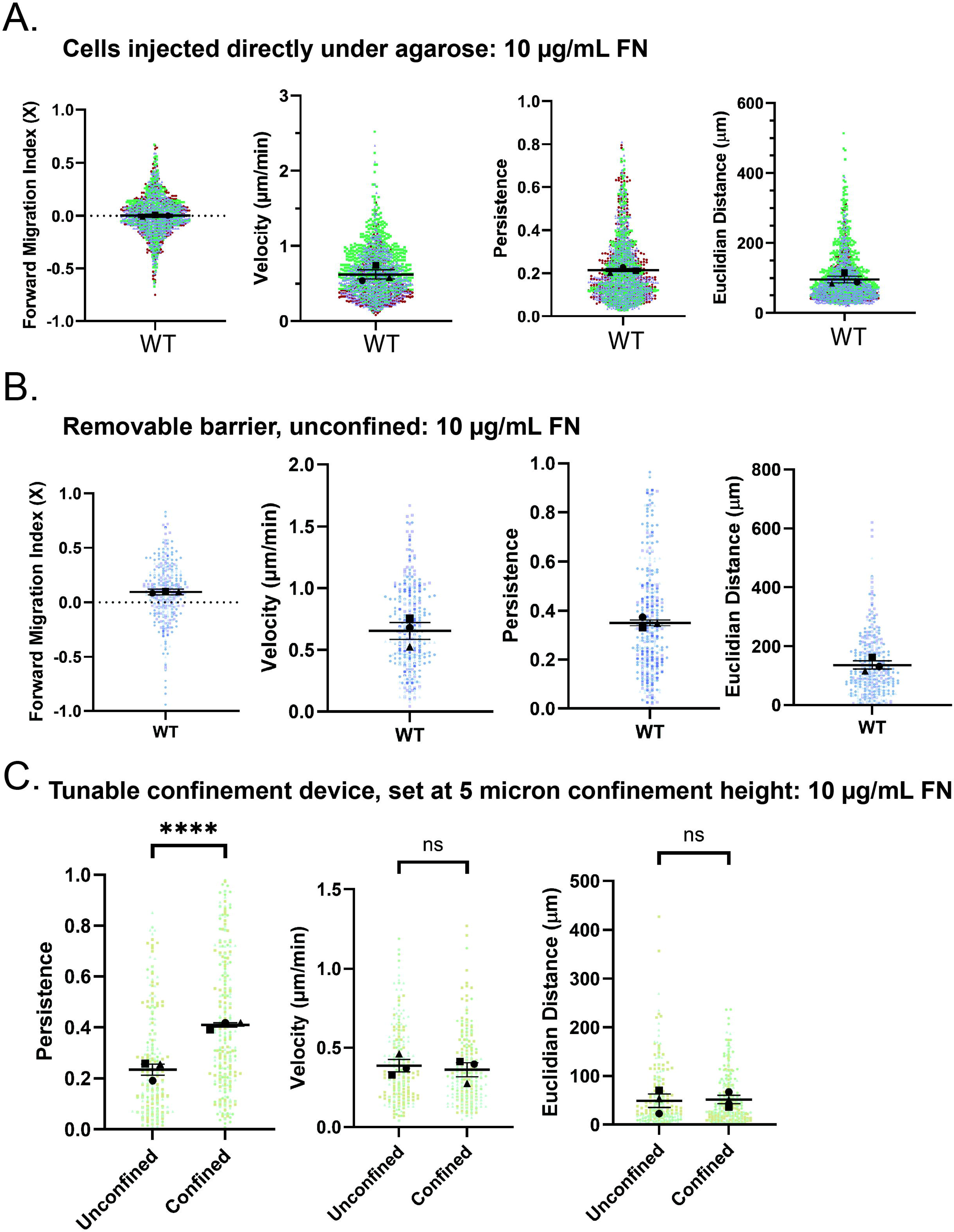
Migration characteristics of cells injected directly under agarose, contained in a removable barrier, or within a tunable confinement device (all on 10 µg/mL fibronectin) (A) FMIx, velocity, persistence and Euclidean distance of cells injected directly under agarose in presence of uniform 10 µg/mL FN. Experimental means and SEM for each experimental replicate are represented with black symbols, and all individual values are plotted and each experimental run is color- and shape-coded (circle, square, triangle). n = 1,591 tracks. These data were pooled from 3 independent experiments. (B) FMIx, velocity, persistence, and Euclidean distance of cells concentrated in a 0.4 mm well within a removable barrier in the presence of uniform 10 µg/mL fibronectin. All cell migration was quantified after the barrier was removed. Experimental means and SEM for each experimental replicate are represented with black symbols, and all individual values are plotted and each experimental run is color- and shape-coded (circle, square, triangle). n = 376 tracks, pooled from 3 independent experiments. (C) Persistence, velocity and Euclidean distance of confined and unconfined cells in the one well static confinement system, set to a confinement height of 5 microns in the presence of uniform 10 µg/mL fibronectin. All cell migration was quantified immediately after confinement was engaged. Experimental means and SEM for each experimental replicate are represented with black symbols, and all individual values are plotted and each experimental run is color- and shape-coded (circle, square, triangle). Statistical analysis was done by Mann-Whitney test. ****p < 0.0001, ns = not significant. n = 248 (unconfined) and 291 (confined) tracks, pooled from 3 independent experiments.

**Supplemental Figure 4.**
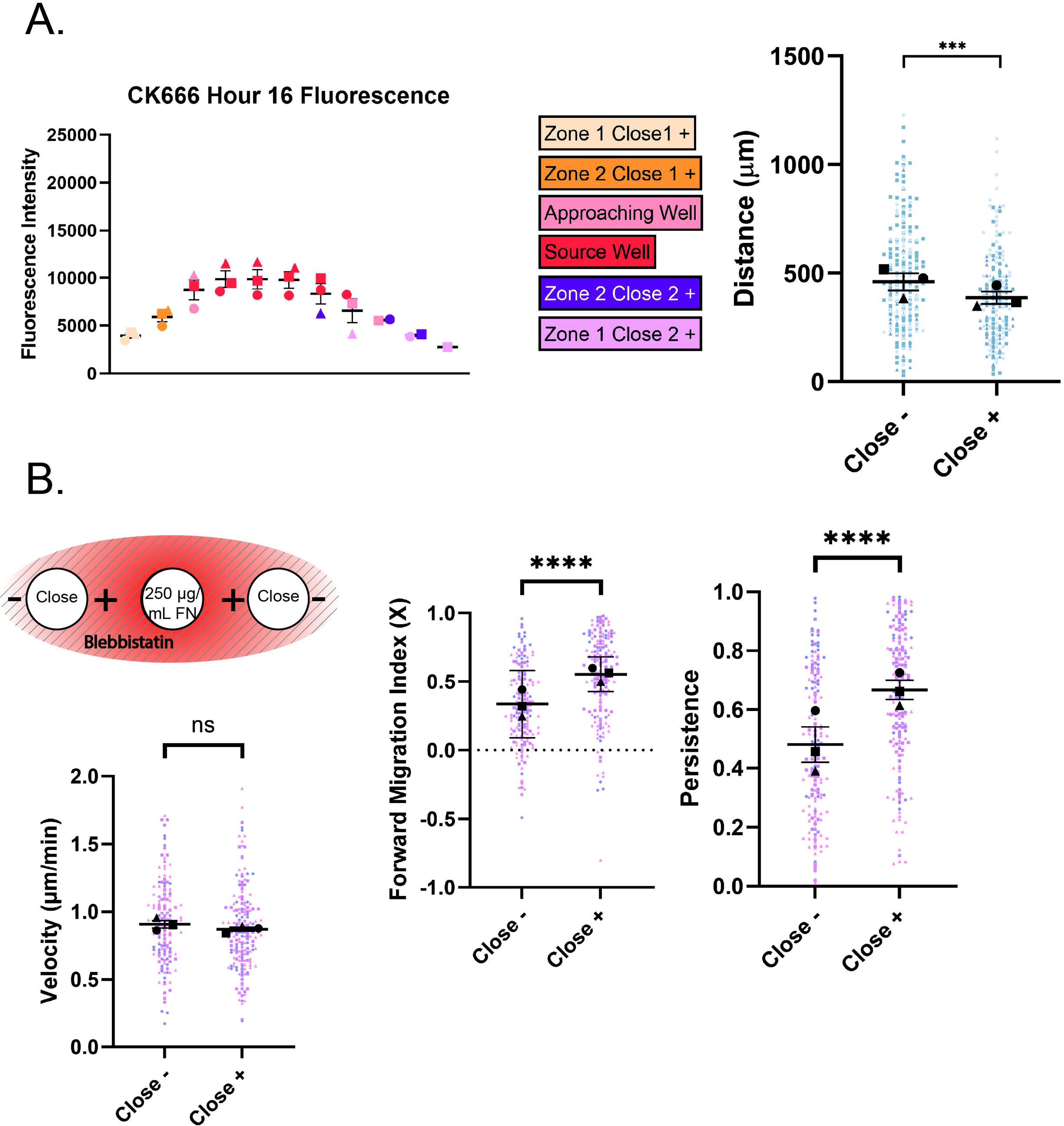
Additional quantification of CK-666 haptotaxis experiment, and blebbistatin haptotaxis results related to Figure 3. (A) *Left*: RRX-FN fluorescence intensity at the experimental endpoint across the well in CK-666 inhibitor experiments. *Right*: Euclidean distance of Close +/- macrophages migrating under agarose in the presence of an RRX-FN gradient and CK-666. Values were calculated from the same dataset as in Fig. 3B. Means and SEM are represented with black symbols, and all tracks are plotted and each experimental run is color- and shape-coded (circle, square, or triangle). Statistical analysis was done by Mann-Whitney test. ***p = 0.0002, ns = not significant. Close-*n* = 250 tracks, Close+ *n* = 194 tracks. Data were pooled from 3 independent experiments. (B) *Left*: Schematic depicting Blebbistatin experimental conditions. A 250 µg/mL FN gradient is generated as normal, but in this case the agarose is made up with 30 µM Blebbistatin, and 15 µM Blebbistatin is included in center well, and both cell wells. Solid lines running through the schematic represent Blebbistatin polymerized into the gel. FMIx, persistence and velocity measurements of macrophages moving under agarose in the presence of blebbistatin. Experimental means from each run are represented with 95% confidence interval (FMIx) or SEM (velocity, persistence), and all individual values are color- and shape-coded (circle, square, triangle). Statistical analysis was done by Mann-Whitney test. ****p < 0.0001, ns = not significant. Close-n = 184 tracks, Close+ n = 200 tracks. Data were pooled from 3 independent experiments.

**Supplemental Figure 5.**
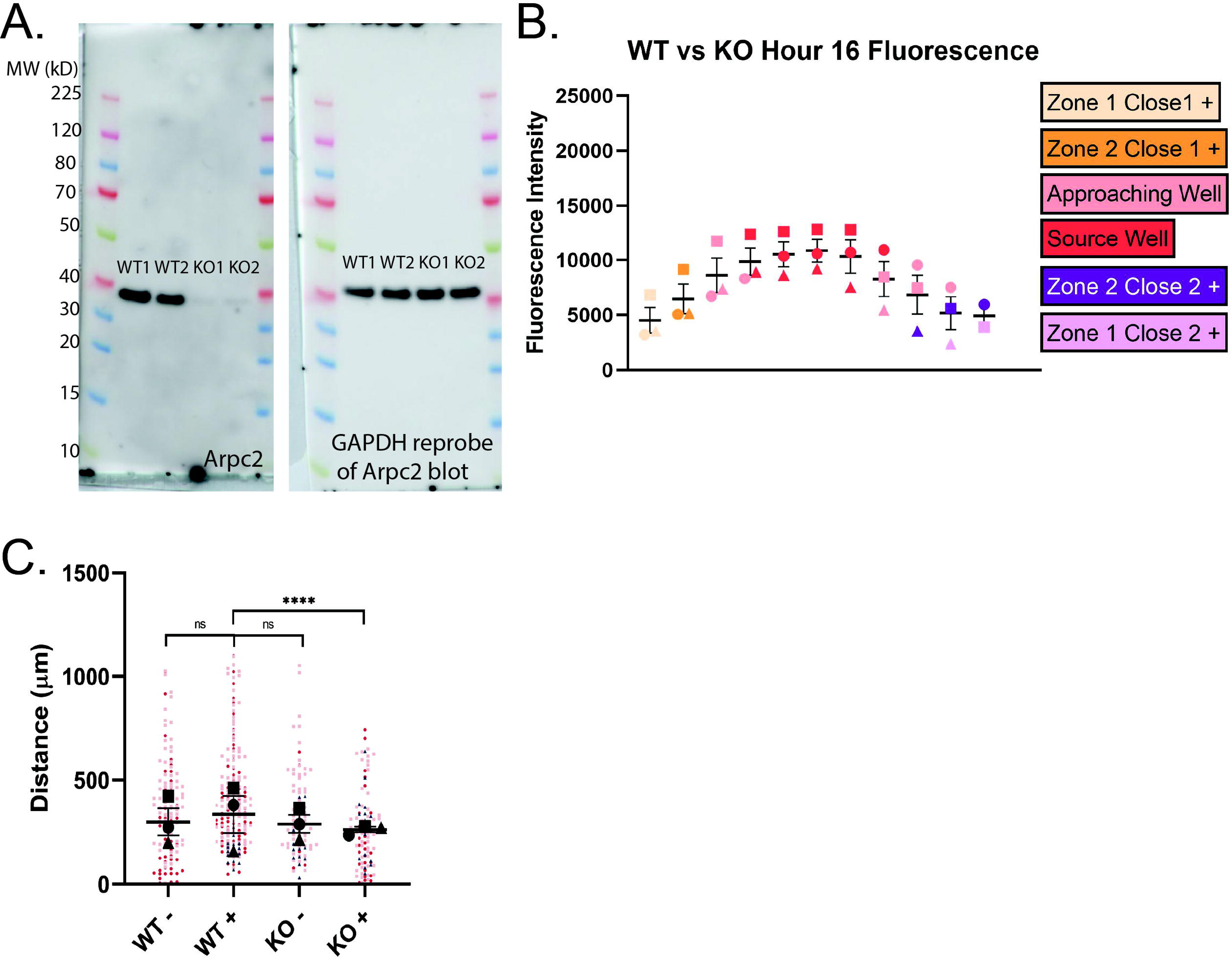
Validation of *Arpc2-/-* macrophages and additional data from WT and *Arpc2-/-* haptotaxis experiment related to Figure 3. (A) *Left*: Overlay of PVDF membrane and western blot of Arpc2. Two populations left untreated (WT) and two populations treated (KO) with tamoxifen to induce tamoxifen-dependent deletion of the Arpc2 subunit of Arp2/3. *Right*: Re-probe of Arpc2 blot for GAPDH to demonstrate equal loading. Molecular weights of markers are indicated to the left of each blot. (B) RRX-FN fluorescence intensity at the experimental endpoint across the well in WT versus KO haptotaxis experiments. (C) Euclidean distance of WT +/- and *Arpc2-/-* (KO) +/- macrophages migrating under agarose in the presence of an RRX-FN gradient. Values were calculated from the same dataset as in Fig. 3D. Means and SEM are represented with black symbols, and all cell migration tracks are plotted and each experimental run is color- and shape-coded (circle, square, or triangle). Statistical analysis was done by Kruskal–Wallis with Dunn multiple comparisons test. ****p < 0.0001, ns = not significant. WT-*n* = 128 tracks, WT+ *n* = 202, KO-*n* = 95, KO+ *n* = 97. Data were pooled from 3 independent experiments.

**Supplemental Figure 6.**
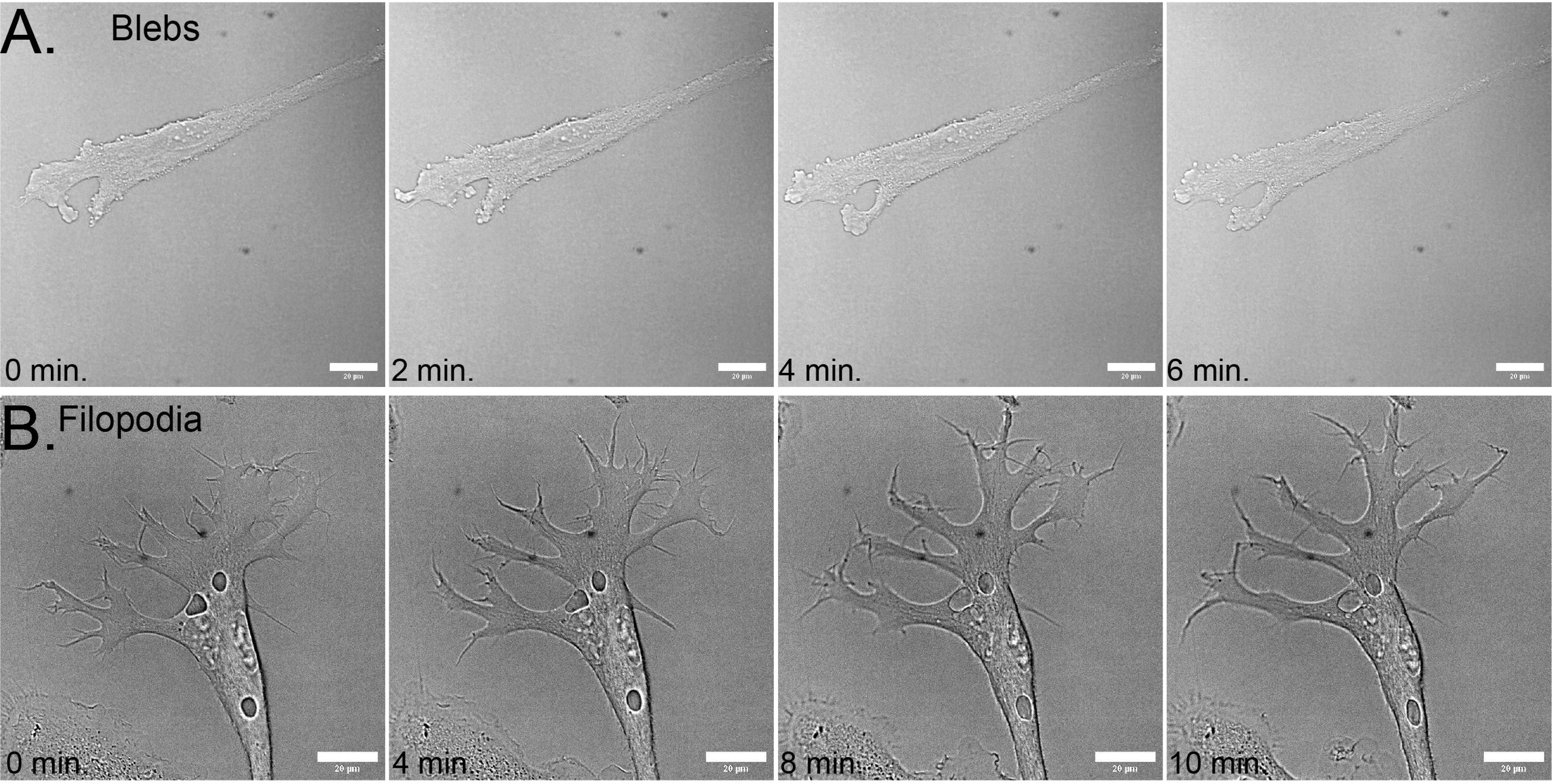
A*r*pc2*-/-* protrusion types. (A) Example images of an *Arpc2-/-* macrophage protruding via membrane blebbing. (B) Example images of an *Arpc2-/-* macrophage protruding via filopodial projections. Scale bar = 20 microns

